# Rapid Estimation of SNP Heritability using Predictive Process approximation in Large scale Cohort Studies

**DOI:** 10.1101/2021.05.12.443931

**Authors:** Souvik Seal, Abhirup Datta, Saonli Basu

## Abstract

With the advent of high throughput genetic data, there have been attempts to estimate heritability from genome-wide SNP data on a cohort of distantly related individuals using linear mixed model (LMM). Fitting such an LMM in a large scale cohort study, however, is tremendously challenging due to its high dimensional linear algebraic operations. In this paper, we propose a new method named PredLMM approximating the aforementioned LMM motivated by the concepts of genetic coalescence and gaussian predictive process. PredLMM has substantially better computational complexity than most of the existing LMM based methods and thus, provides a fast alternative for estimating heritability in large scale cohort studies. Theoretically, we show that under a model of genetic coalescence, the limiting form of our approximation is the celebrated predictive process approximation of large gaussian process likelihoods that has well-established accuracy standards. We illustrate our approach with extensive simulation studies and use it to estimate the heritability of multiple quantitative traits from the UK Biobank cohort.

## 1 Introduction

In the past few decades, genome-wide association studies (GWASs) have identified hundreds of single nucleotide polymorphisms (SNPs) influencing the genetic architecture of complex diseases and traits. For majority of the traits, however, the associated SNPs from a GWAS only explain a small fraction of the heritability estimated using twin and family studies. In search of this so called “missing heritability”, there were attempts to capture even infinitesimal SNP effects by taking into account genome-wide variants in a linear mixed model (LMM) framework [32, 19, 22, 8]. The SNP-based LMM framework, often known as genome-based restricted maximum likelihood (GREML) approach, usually involves distantly related people, whose extent of genetic relatedness depend on their evolutionary history [31]. The total trait variance in this LMM approach is decomposed into two variance components such as the additive genetic variance and the residual variance. [24, 28]. The approach requires computation and inversion of a high-dimensional genetic relationship matrix (GRM) from the genome-wide SNP data of dimensionality same as the sample size. Heritability is calculated as the ratio of the additive genetic variance to the total variance and the parameters are usually estimated using a restricted maximum likelihood (REML) approach.

In recent years, advances in genome sequencing have generated huge amount of genetic data on large scale cohort studies, such as UK Biobank [1], Precision Medicine cohort [17], Million Veterans Program [13]. These studies collect data on millions of genetic markers and numerous diseases/traits on thousands of individuals. For example, UK Biobank cohort has data on approximately 500,000 individuals, 800,000 markers and numerous traits. Therefore, it is needless to say that GREML based methods need to be extremely time and memory efficient to be applicable on such magnanimous studies.

Programs such as genome-wide complex trait analysis (GCTA) [32], genome-wide efficient mixed model association (GEMMA) [34] have implemented efficient algorithms to fit the GREML approach. These programs usually follow two steps: first, compute the genetic relationship matrix (GRM) with the SNP data on the individuals and second, use the computed GRM to fit a GREML corresponding to a trait. If *N* be the number of individuals and *M* be the number of SNPs, the first step of computing the GRM, takes complexity of *O*(*MN* ^2^) FLOPS (floating point operations). And, the next step i.e fitting the GREML to estimate heritability, requires inverting the GRM matrix which uses per iteration complexity of *O*(*N* ^3^) FLOPS. When *N* is extremely large (say more than 100,000), this step becomes computationally intractable. It should be noted that the first step (computing the GRM) is also computationally very demanding (especially when *M, N* both are large). In large biobank-scale studies, where the interest is to estimate heritability of a large number of traits, implementing these approaches become computationally very demanding.

Recently, an approximate method named Bolt-REML [20, 21, 22] has been proposed that trades off small amount of accuracy in favor of greater computational speed. It follows a different path than the above methods. It does not compute the GRM but uses the SNP data directly to fit the GREML by monte carlo average information REML algorithm. It has computational complexity of *O*(*MN* ^1.5^) per iteration which is better than the previous methods in terms of *N*. The software is well optimized and in our analysis of UK Biobank data, it performed much better compared to the other approaches in terms of computational time. However, the complexity of Bolt-REML is not linear in *N* which makes it challenging to use for larger *N* (> 300, 000). Additionally, the computational complexity also increases linearly with *M*. Thus, in a large cohort with millions of SNPs, it would be immensely intensive to use Bolt-REML for estimating heritability of all the traits one by one. On the other hand, the previous approaches estimate the GRM only once (that requires computational complexity of *O*(*MN* ^2^)) and after that, the complexity of analyzing any trait does not depend on *M*.

In this paper, we approximate the likelihood of the GREML approach to develop a rapid algorithm for estimating heritability. The approximation is motivated by the concepts of genetic coalescence [18, 9] and gaussian predictive process models [3, 11]. Our proposed approach PredLMM exploits the structure of the GRM to ease the computationally demanding linear algebraic steps of the standard GREML algorithm such as calculation of the determinant or inverse of a high dimensional matrix (*N × N*) at every iteration. It reduces per iteration computational complexity from *O*(*N* ^3^) FLOPS (floating point operations) to *O*(*Nr*^2^)+ *O*(*r*^3^) FLOPS where *r* is much smaller than *N*. Theoretically, we show that under a model of genetic coalescence, the limiting form of our approximation is the celebrated predictive process approximation of large gaussian process likelihoods [3] that has well-established accuracy standards. Empirically we have verified the reliability and robustness of our proposed approach through extensive simulation studies by replicating many possible realistic scenarios. We have analyzed the UK Biobank cohort data (with 286,000 British individuals and 566,000 SNPs) to estimate the heritability of *Standing Height, Weight, BMI, Systolic and Diastolic blood pressure, Hip and Waist circumference*. We have implemented PredLMM in an efficient Python module (https://github.com/sealx017/PredLMM).

## 2 Materials and Methods

### 2.1 Genome-based Restricted Maximum Likelihood

#### 2.1.1 Model Specification

Let **Y** denote the *N ×* 1 vector of phenotype corresponding to *N* individuals, **X** denote the *N × p* matrix of covariates, and **W** denote the *N × M* matrix of mean and variance scaled genotype for the *N* individuals and *M* SNPs, i.e., *E*(*w*_*ij*_) = 0 and *V ar*(*w*_*ij*_) = 1. Consider the following LMM,

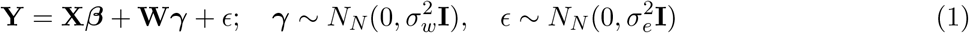

And, the corresponding marginal model can be written as,

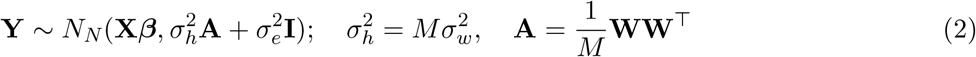

where **A** is formally known as the Genetic Relationship Matrix (GRM) and **I** is the identity matrix. Heritability is calculated as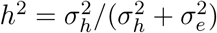.

#### 2.1.2 Estimation Approaches

To estimate the variance parameters 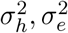, and eventually *h*^2^, different REML algorithms are generally used. Thus, the entire framework is referred to as genome-based restricted maximum likelihood (GREML) approach. There are two types of programs implementing the GREML approach: a) Exact Methods (methods that converge to the REML optimum) and b) Approximate Methods (methods that approximate the REML optimum).

### Exact Methods

Programs such as GCTA [32], GEMMA [35] operate in two steps: first, compute the GRM, 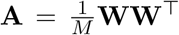 and second, consider the computed **A** in the marginal model from Equation to estimate *h*^2^ using different REML algorithms. These REML algorithms are iterative and compute analytically exact solutions. For example, GEMMA uses a modified version of Newton-Raphson method (considers exact Hessian), GCTA uses average information (AI) method (considers approximate Hessian and hence, computationally faster). The second step involves computing the inverse and determinant of the *N × N* dense matrix 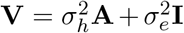 at every iteration which takes *O*(*N*^3^) FLOPS, making these exact methods computationally intractable as *N* increases.

### Approximate Methods

Unlike the previous methods, Bolt-REML [20, 21, 22] does not compute the GRM **A**. It directly uses with the SNP data matrix **W** and follows a Monte Carlo REML approach that uses random sampling to approximate the derivatives of the log likelihood corresponding to the marginal model from Equation 2. The algorithm has computational complexity of *O*(*MN* ^1.5^) per iteration which is better than the previous methods in terms of *N*. The software is well optimized and in our analysis of UK Biobank data, unlike the previous methods, it would successfully converge for moderately large *N* (*N >* 100, 000) in a reasonable amount of time. However, the per iteration computational complexity of Bolt-REML still increases linearly with *M*. Thus, in a cohort study where *M* is closer to a million (or, larger), it will become computationally much more challenging to use Bolt-REML. On the other hand, the previous approaches estimate the GRM only once (with computational complexity of *O*(*MN*^2^)) and after that, the complexity of analyzing any trait does not depend on *M*.

### 2.2 Sub-sample based GREML

Since, the likelihood based methods above involving the full population become increasingly computationally demanding as the population size *N* increases, an alternative would be to utilize a sub-sample based approach described here:

- Choose a random sub-sample *k* of size *n*^*\**^ from the dataset such that a standard GREML based program, for example GCTA can be used to estimate heritability 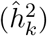.
- Repeat the step above *K* times and take the average of the heritability estimates: 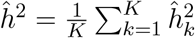 as an estimate of *h*^2^.

Theoretically, the final estimate of heritability: 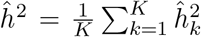 should be consistent but with a much higher variance than the full data based GREML estimate. Also, it would be challenging to obtain a closed form expression of the variance (unless *K* = 1) and one would have to resort to bootstrapping for that purpose. Additionally, though this approach can drastically reduce the computational complexity, choosing the sub-sample size: *r* and the number of repetition: *K* to achieve a reliable estimate of heritability may become an arduous task itself. In our simulation studies and real data analysis, we assess the performance of a simple version of this method by considering a single repetition and varying the value of *n*^*\**^. We refer to it as GREML (sub).

### 2.3 Proposed Method

#### 2.3.1 Asymptotic limit of the GRM

First, we show that under certain assumptions, as the number of SNPs *M* goes to infinity, the likelihood corresponding to the marginal model from (2) converges almost-surely to a gaussian process (GP) likelihood. The assumptions are as follows,

1. *Assumption 1 (Correlation across individuals):* We assume that each individual *i* = 1, 2, …, *N* can be represented by a point (location) **s**_*i*_ in an abstract spatial manifold 𝒟 equipped with a distance *d*. The correlation between the genotypes of individuals *i* and *i*′ at the *j*^*th*^ SNP is given by *Cov*(*w*_*ij*_, *w*_*i′j*_) = *C*_*j*_(**s**_*i*_, **s**_*i*_′) where *C*_*j*_ is a valid covariance function in *D* which decreases monotonically with increasing distance 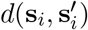. This assumption is rooted in the theory of *genetic coalescence* [9, 18]. The coalescence model describes the relationships within a sample from the present individuals (sequences) back to the most recent common ancestor (MRCA) [30]. Under coalescence, the correlation between genotypes of individuals will vary inversely with the *time to coalescence*, i.e., number of ancestral generations till the *most recent common ancestor*. Hence, the MRCAs of different pairs of individuals in a sample can be assigned to nodes of a genealogical tree. Trees are equipped with a valid distance metric (shortest distance between nodes) and models for tree-structured objects commonly specify the correlation as decreasing function of the distance [4]. The maximum likelihood estimate of *h*^2^ from (2) has been shown to be consistent in [16]. The theory relies on the assumption that the genotype distributions are independent across individuals (upto standardization). Formally, **w**_*i*_ ⊥ **w**_*i′*_ for any two individuals *i*  *i*′ where **w**_*i*_ = *i*^*th*^ row of **W**, is the genotype vector for the *i*^*th*^ individual. Such an assumption of between-individual independence of genotype distributions is in sharp violation of the principles of coalescent theory. We note while coalescence model is a natural example where our assumption of latent embedding is realized, the concept is not just restricted to trees and can be compatible with more complex models of ancestry depicted by any manifold with a notion of distance.
2. *Assumption 2 (Stationarity and ergodicity across the SNPs):* We assume that the centered and scaled genotype process 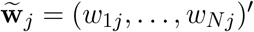 is second-order stationary and ergodic for *j* = 1, 2, .... Stationarity translates to 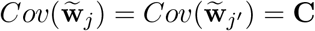 for all *j, j*′ implying that the covariance functions *C*_*j*_ = *C* for all *j* = 1, 2, … Ergodicity implies that as the number of SNPs grows, we have.

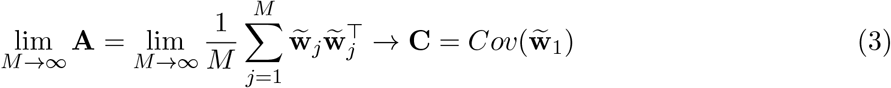

The simplest setting where this assumption is satisfied is when the scaled and centered genotype processes 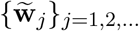 are assumed to be iid. Assumption of iid genotypes is common in theoretical studies of the heritability estimation [16] but independence is only sufficient and not necessary for us. More realistic scenarios like presence of linkage disequilibrium (LD) that effectuates correlation across genotypes can also be accommodated as long as the ergodicity is ensured. As shown in [27], the pairwise LD among loci in a homogeneous population decreases exponentially as a function of the genetic distance, which validates the feasibility of our assumption. Correlation structures arising from absolutely regular-mixing processes [6] like autoregressive (*AR*(*p*)), moving average (*MA*(*q*)) or *ARMA*(*p, q*) [23] will satisfy the strong law of convergence in Equation (3) [25].

Under Assumption 2, we have the following assertion on the limit of the marginal LMM likelihood from

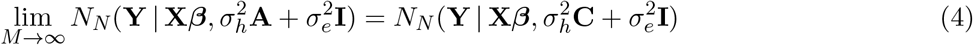

where *N*_*N*_ (**Y** | ***µ*, Σ**) denotes the *N*-variate normal likelihood for a realization **Y** with mean ***µ*** and variance **Σ**. Thus the likelihood used in heritability estimation converges to a likelihood for data **Y** a partial realization of a Gaussian process on 𝒟 with mean 0 and covariance function *C* observed at the *N* latent locations **s**_1_, **s**_2_, …, **s**_*N*_. It is expected that estimation of heritability using the limiting likelihood (4) will be similar to that from the exact likelihood (2) as the number of SNPs *M* is usually very large.

#### 2.3.2 PredLMM

Just switching to the limiting likelihood (4) does not ease any of the computational burden as GP likelihoods also require *O*(*N*^3^) FLOPS. However, over the last two decades a series of increasingly sophisticated algorithms have been proposed for fast approximate GP likelihoods [see 15, for a recent review].

Our approach uses predictive process (PP) [3, 11] which results in the low-rank plus diagonal approximation of the dense matrix **C**. Let 𝒮 = {**s**_1_, **s**_2_, …, **s**_*N*_} denote the set of *N* latent locations, and 𝒮* = {**s**_1_, **s**_2_, …, **s**_*r*_} denote a set of *r ≪ N* locations in 𝒟 referred to as the *knots*. Also, for two sets *A* and *B* in 𝒟 let **C**_*A,B*_ denote the |*A*|*×*|*B*| matrix (*C*(**s**_*i*_, **s**_*i′*_))_*i∈A,i*_′∈*B*. The predictive process approximation of **C** is given by

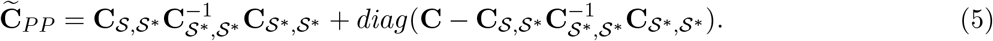

The first term is a low-rank factorization as the number of knots is much less than the sample size. [3] showed that this low-rank term is the optimal (in terms of reverse Kullback Leibler divergence) low-rank approximation of **C** using the knots **S**^*\**^. [11] proposed adding the diagonal matrix (second term) to eliminate a positive bias on the diagonal entries. For moderate choices of *r ≪ N*, inference from the predictive process likelihood provides an excellent approximation to that from the full GP likelihood. Computationally, predictive process only requires *O*(*Nr*^2^ + *r*^3^) FLOPS and as *r ≪ N*, the approximation results in massive gains in run times. Consequently, predictive processes is one of the most popular approximations of the full GP likelihood and is widely adopted in many spatial applications.

In our setting, direct usage of predictive process likelihood is not recommended for two reasons. First, the locations **s**_*i*_ are unknown to us. Hence, **C**_*PP*_ can only be calculated using approximate locations like a vector of the top few PC scores. The impact of such choices of locations is less clear. Second, covariance functions usually involve additional spatial parameters ***θ***, thereby increasing the number of unknown parameters to be estimated.

Instead, we consider the following strategy. We choose 𝒮* to be a subset of 𝒮, and define ℐ to be the subset of *ℬ* = {1, 2, …, *N*} containing the indices corresponding to *S\**. We can decompose the GRM **A** as,

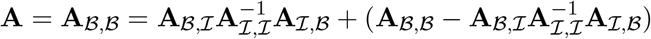

The decomposition is inspired by the concept of conditional variance [10]. The first term 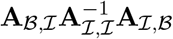 on the right is the low-rank part of the full GRM **A** that is explained by the information about the subset of individuals ℐ, while the second term 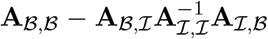 is the residual GRM of the individuals in the subset ℬ ∩ ℐ^*c*^ that is not explained by the individuals in the subset ℐ. Replacing the term on the right with its diagonal, we then have a direct low-rank plus diagonal approximation of **A** as

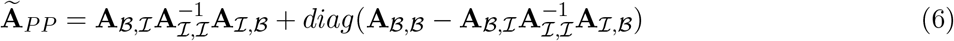

We propose using the likelihood 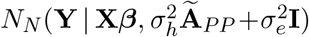 for heritability estimation. It is clear that **A** and **Ã**_*PP*_ agree on the diagonals, and on the sub-matrix corresponding to the knots ℐ. Also, lim_*M*→∞_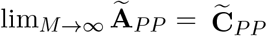. Hence, using triangular inequality, we can write

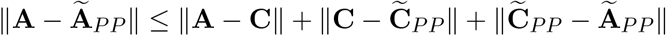

Under assumption 2, the first and third terms vanishes as *M* → ∞, while for a well chosen set of knots **S**^*\**^, the predictive process approximation 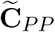 is close to **C** (since *C* is a decreasing function of the distance as postulated in Assumption 1). Hence the middle term will also be small. This justifies why for large *M*, **Ã** _*PP*_ is expected to be close to **A**.

In our empirical studies detailed later, the predictive process approximation consistently and substantially outperforms the subsample-based method when both uses the same set of knots (subsample) ℐ. We offer some insight into this. The first term of **Ã**_*PP*_ in Equation 6 is

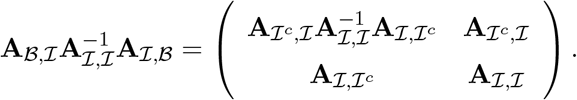

As mentioned before, this low-rank matrix is the best estimate of **A** based on the genetic information only from the individuals in subset ℐ and their genetic correlation with the individuals in subset ℐ*c*. If using the subsampling based approach with the same subsample ℐ, one would only use the submatrix **A**_ℐ,ℐ_ to estimate *h*^2^. This thus ignores the genetic correlation of these subsampled inviduals with those not subsampled (quantified as **A**_ℐ,ℐ_*c*), and is thus suboptimal to the predictive process approach which leverages this genetic relationship among individuals while remaining computationally scalable.

#### 2.3.3 Computational gains

Evaluation of our PredLMM likelihood 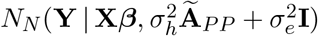, does not require computing or storing the entire *N* × *N* GRM matrix **A** and can be calculated only using the *N* × *r* tall thin sub-matrix **A**_*ℬ,ℐ*_, the small *r* × *r* square matrix **A**_*ℬ,ℬ*_, and diagonal elements of **A**. This reduces storage from *O*(*N* ^2^) to *O*(*Nr* + *r*^2^) – a substantial gain for biobank-scale studies with large *N* as *r ≪ N*.

Subsequently, the nice low-rank plus diagonal structure of **Ã**_*PP*_ facilitates fast evaluation of the likelihood. Inverse of 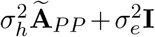 becomes feasible and significantly rapid using the Woodbury matrix identity [29], while the matrix determinant lemma [14] is leveraged for scalable computation of the determinant. Both the steps involve *O*(*Nr*^2^ + *r*^3^) FLOPS, as *r ≪ N*, the computation is thus becomes linear in *N* – a drastic reduction from the *O*(*N*^3^) FLOPS required for evaluating the true likelihood.

#### 2.3.4 Choice of knots design and number

In traditional applications of gaussian processes in spatial statistics, the domain 𝒟 is known and the locations **s**_*i*_ are observed. Hence, the knots need not coincide with the data locations. Recommended choices for the knot-set include space-filling designs and lattices [3]. In our case, the locations are artificial constructs to motivate our direct approximation. Hence, restricting the knot set to be a sub-sample of these hypothetical data locations is necessary to ensure that the direct approximation **Ã**_*PP*_ can be calculated using submatrices of **A**. However, our practice has precedence even in conventional spatial settings. Using some of the data locations has been shown to improve performance of predictive process [3], while related approaches like splines and other basis function expansions also commonly use data locations as knots. We used random sub-samples as knots, as empirical experiments highlighted in Section 3 demonstrated considerable robustness of results to the choice of sub-sample. More informed choices for knots like cluster representatives from PCA-, GRM-, or dendogram-based clustering can also be used.

Choice of the the number of sub-samples *r* to be used for PredLMM is more nuanced. Performance of predictive process is generally more sensitive to the number than the design of the knots [3]. Increasing *r* improves the quality of the approximation, with **Ã**_*PP*_ exactly equalling **A** when *r* = *N* and ℐ = ℬ. However, as the computation is cubic in *r*, use of a very large *r* would defeat the purpose of the approximation. In practice, estimating *h*^2^ for a few choices of *r* of increasing values is recommended, stopping when the estimates of *h*^2^ no longer change substantially. Parallel computing resources, if available, can be heavily deployed for this step.

#### 2.3.5 Asymptotic variance of the estimator

We have derived the expression of the asymptotic variance (standard error) of the PredLMM estimator. Since it is extremely time consuming to perform the matrix multiplications needed for the exact computation of the variance expression, we make some reasonable approximations. The details of the derivation can be found in Appendix A.1.

## 3 Results

### 3.1 Simulation Study 1: Simulation under Coalescent Model

The following simulation study replicates a scenario when Assumption 1 from Section 2.3.1 approximately holds i.e every individual originates from a common ancestor and individuals in the same sub-population share a more recent ancestor than the individuals in different sub-populations. A tree structure with four generations has been depicted in Figure 1.

**Figure 1:**
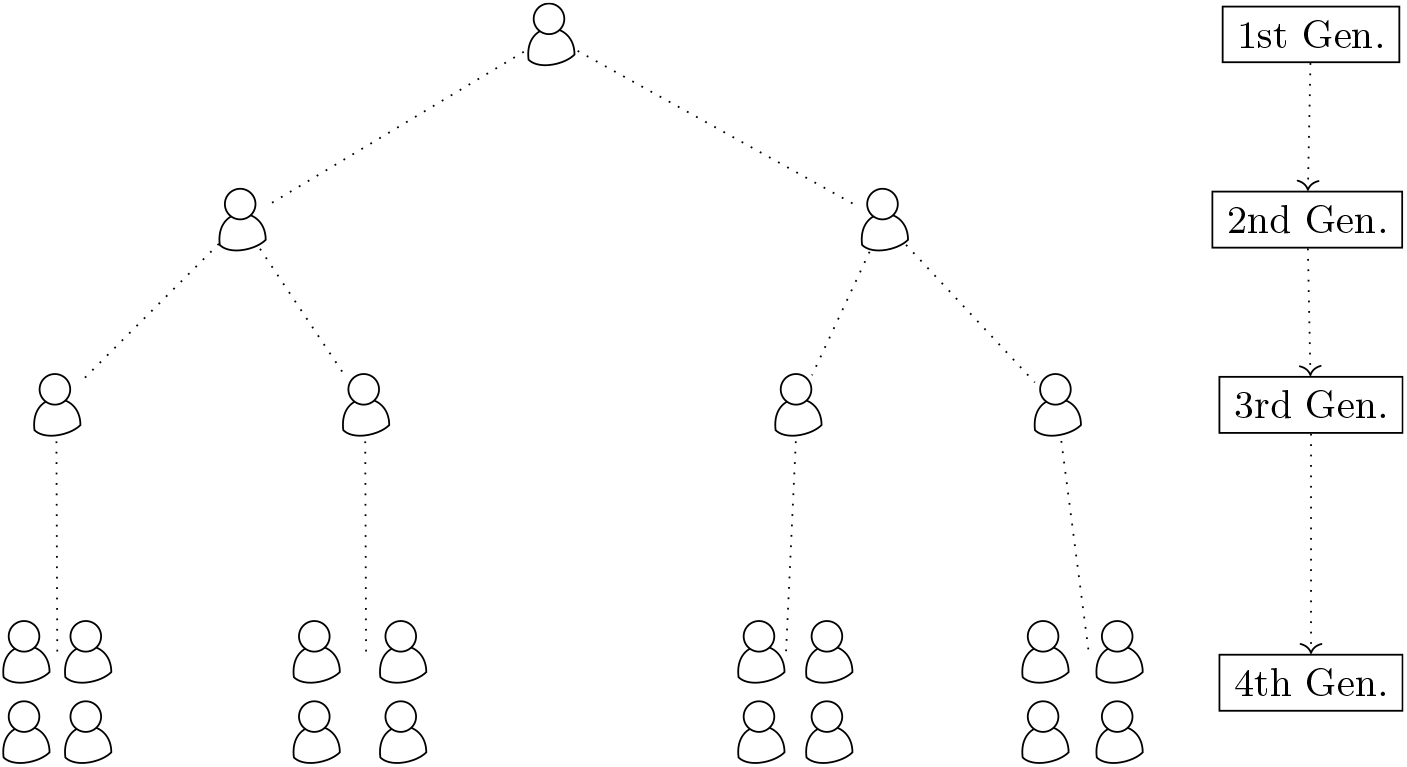
The figure shows a model coalescence with four generations. Each of the four blocks of people in the fourth generation share one of the individuals from the third generation as common ancestor. Similarly, the four people in the third generation have originated from the two in the second generation. And, finally those two people have originated from a common ancestor in the first generation.

As in Figure 1, we generated a population from a coalescence model. For each SNP *j* (*j* = 1, …, *M*), the allele frequency 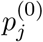 in the first generation was drawn from a uniform distribution on [0.1, 0.9]. In the second generation, allele frequencies of two different individuals: 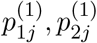 were independently simulated from a beta distribution with parameters 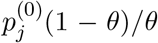 and 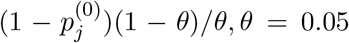. This model is commonly known as Balding-Nichols model [2, 26]. In the third generation, allele frequencies of two individuals: 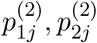 were independently drawn from a beta distribution with parameters 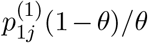 and 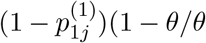 and allele frequencies of other two individuals: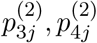 were independently drawn from a beta distribution with parameters 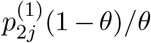 and 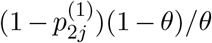. Finally, in the fourth generation, the allele frequency of *j*-th SNP of the *i*-th individual from the *k*-th sub-population (*k* = 1, …, 4): *p*_*ijk*_ was generated from a beta distribution with parameters 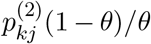 and 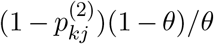. We kept the size of each of the four sub-populations at *N*/4 so that the total population size is *N*. We next simulated the SNP genotype: *w*_*ijk*_ from a binomial distribution: *Bin*(2, *p*_*ijk*_) assuming Hardy-Weinberg equilibrium. Once we simulated genetic data for *M* markers on *N* individuals with **W** representing a *N* × *M* matrix, we randomly selected *m*_*causal*_ many causal SNPs (columns) to create a *N × m*_*causal*_ dimensional matrix **W**^*causal*^ which were then used to simulate the phenotype. Fixed effect for each of the causal SNPs: *u*_*m*_was simulated from *N* (0, *h*^2^*/m*_*causal*_), and the residual effect *e* was simulated from *N*_*N*_ (**0**, (1*/h*^2^ − 1)**I**_*N*_). Finally, the phenotype vector was generated as, 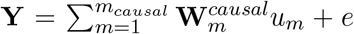 with 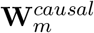 as the *m*-th column of **W**^*causal*^.

We considered two different values of the true heritability: *h*^2^ (low and high) and two different combinations of the number of individuals *N* and the number of SNPs *M*. We considered case (1.1): *h*^2^ = 0.2, *N* = 5000, *M* = 8000, case (1.2): *h*^2^ = 0.2, *N* = 8000, *M* = 13000, case (2.1): *h*^2^ = 0.8, *N* = 5000, *M* = 8000 and case (2.2): *h*^2^ = 0.8, *N* = 8000, *M* = 13000 to study the influence of *M* and *N* on heritability estimation. Figure 2 shows the box-plots of the heritability estimates by different methods over 100 repetitions. We considered several full likelihood based GREML methods as discussed in Section 2.1.2: GCTA, GEMMA and Bolt-REML for comparison with PredLMM. Since all of these methods maximize the full likelihood corresponding to the marginal model in (2), their estimates are expected to be precise and quite similar to each other. Figure 2 shows very similar performance of these full likelihood based GREML methods. GREML (500) and GREML (2000) refer to the sub-sample based GREML discussed in Section 2.2 with sub-sample sizes of 500 and 2000 respectively. PredLMM (500) and PredLMM (2000) refers to fitting PredLMM with knot-sizes (*r*) 500, 2000 respectively. Both PredLMM (500) and PredLMM (2000) produced nearly unbiased estimates in all scenarios with PredLMM (2000) giving more precise estimates of heritability. GREML (500) mostly underestimated heritability and had a very large variance in all the cases. The variance was higher for low value of heritability. Our findings show that the sub-sample based GREML can be unreliable if the sub-sample size is small. GREML (2000) performed better with no apparent bias but still had much larger variance than both PredLMM (500) and PredLMM (2000). Overall, we found that PredLMM produced robust estimates of heritability even with a very small set of knots, when the genetic data was simulated from a Balding-Nichols model [2, 26]..

**Figure 2:**
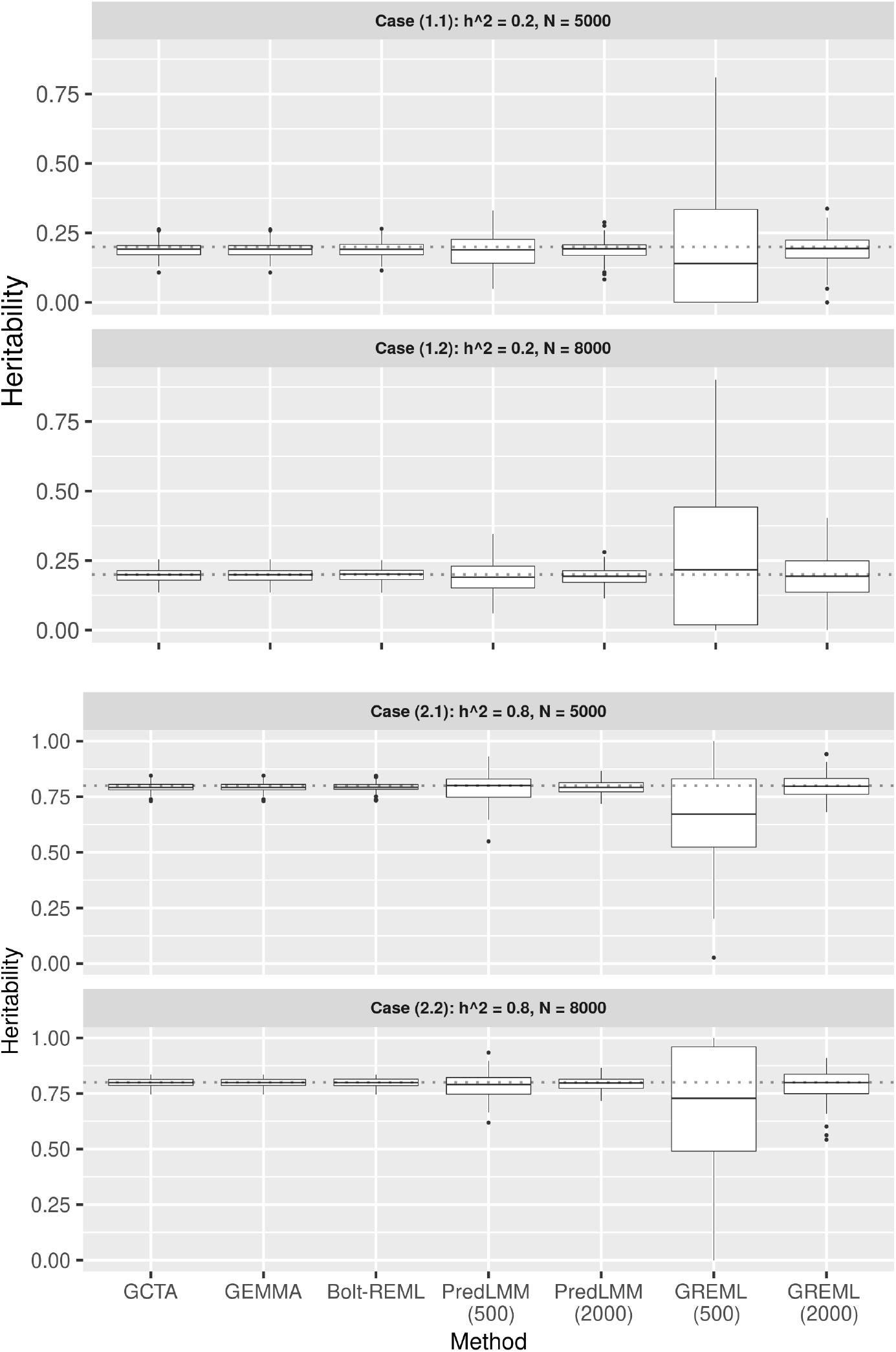
The figure shows the boxplots of the heritability estimates by different methods from Simulation Study 1 (Section 3.1). Each of the four sub-plots corresponds to four different cases. The true heritability value for each case is mentioned in the sub-plot header and is also depicted by the dotted line.

### 3.2 Simulation Study 2: Simulation using UK Biobank data

To replicate more realistic scenarios, we next considered simulations using the UK Biobank cohort data [1]. UK Biobank is a large long-term biobank study in the United Kingdom which is investigating the respective contributions of genetic predisposition and environmental exposure to the development of various diseases. We had access to 784,256 markers and multiple phenotypes on 502,628 individuals. The population is predominantly British (442,687) with a few other ethnicities such as Irish (13,213), Other White (16,340), Asian (9839), and Black (8038). There is clear genetic clustering in the UK Biobank sample that as been explored in [12].

Keeping the genetic heterogeneity in mind, we looked into a diverse group of simulation setup using the genetic data from the UK Biobank study. After standard quality control steps as advised in [7] (removing SNPs with MAF less than 0.01 and missingness over 10%, removing individuals with high missing genotype rate), we had approximately 320,000 individuals and 566,000 SNPs. Since, conducting simulation studies with the entire dataset would be very computationally expensive, we created a sub-population for our simulation study. We randomly selected 157,000 people (120,000 British and 37,000 from other ancestries such as Asian, Black, Irish, and Indians). Majority of the full GRMEL-based methods such as GCTA, GEMMA are computationally infeasible for such a large number of individuals. Bolt-REML is the only full GREML-based method that is still viable in this context. But, as we see from Figure 3 that even for a single simulation with 100,000 individuals, Bolt-REML took approximately 1000 minutes to run (more details regarding the time comparison can be found in Section 4) Thus, we only compared PredLMM with GREML (sub) in the following simulations.

**Figure 3:**
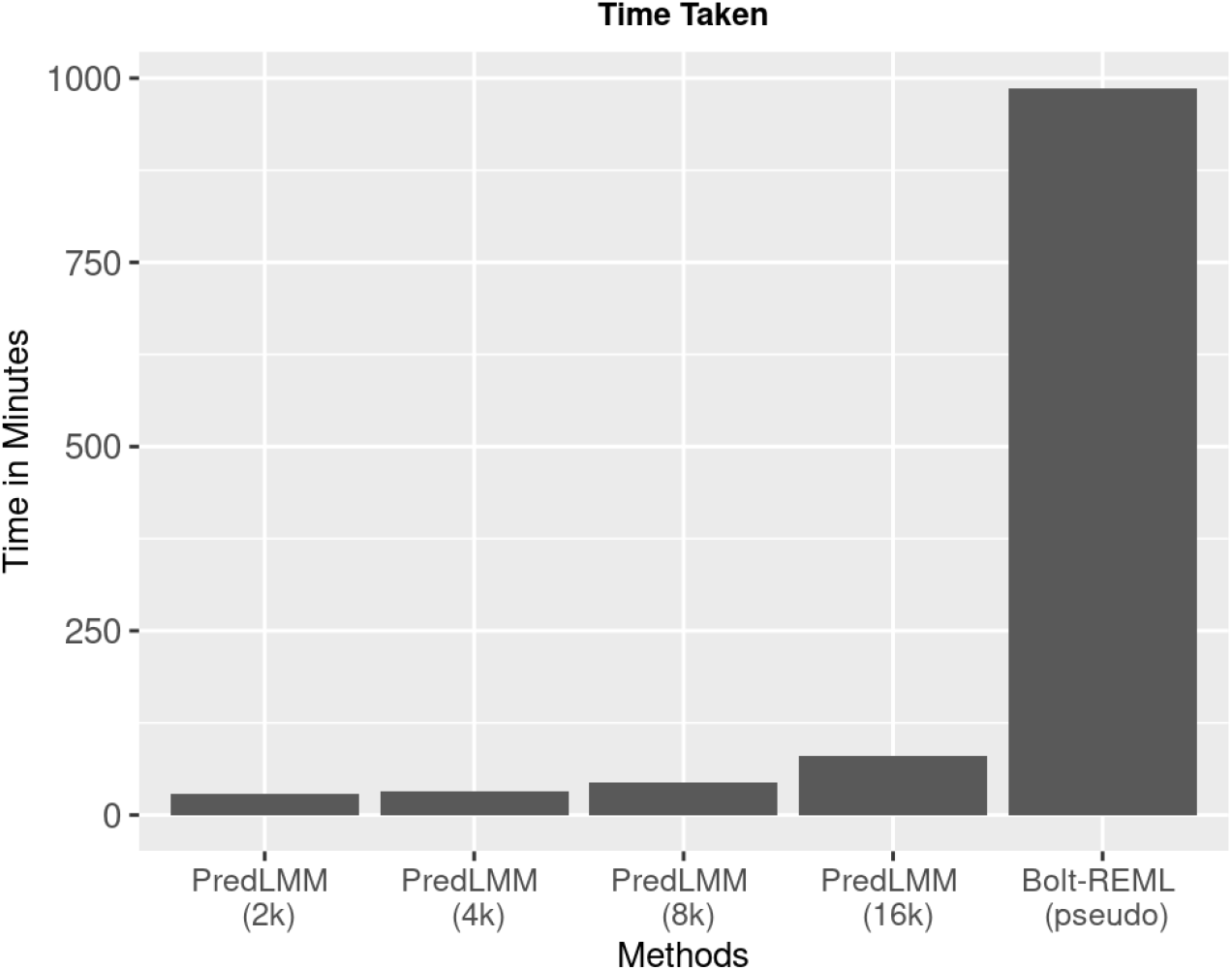
The figure shows the time taken by PredLMM with different knot-sizes such as, 2000, 4000, 8000 and 16,000 and by Bolt-REML for a single simulation with 100,000 individuals and 566,000 SNPs (from Simulation Study 2A).

In our Simulation Study 1, the GREML (sub) appoach produced highly biased estimates. We found that the bias was significantly lower if the population was heterogeneous (with multiple ethnicities). That is why, we considered two sets of simulation below: (2A): one with a homogeneous sub-population (only British ancestry) and (2B): the other with a heterogeneous sub-population of these 157,000 people.

#### 3.2.1 Simulation Study 2A: Simulation with a homogeneous sub-population

We considered three different true values of heritability (low to high): (a) *h*^2^ = 0.2, (b) *h*^2^ = 0.4 and (c) *h*^2^ = 0.6. Every time we randomly selected 100,000 people with British ancestry from the sample of 157,000 people. Next, we simulated the trait as **Y**_100,000_ ∼ *N*_100,000_(**0**, 250*h*^2^**A**_100,000_ + 250(1 *− h*^2^)**I**_100,000_) where **A**_100,000_ was the corresponding GRM. We compared PredLMM with GREML (sub) for four different sub-sample (knot) sizes, 2000, 4000, 8000, 16000. Box-plot comparison of the estimates are shown in Figure 3. PredLMM for small knot-sizes such as 2000, 4000 showed a slight upward bias when *h*^2^ was low [case (a)] and a slight downward bias when *h*^2^ is high [case (c)]. The bias vanished as the knot-size increased. For moderate value of *h*^2^ i.e in case (b), the bias was imperceptible even for the smallest knot-size of 2000. The GREML (sub) estimates however were completely unreliable for small sub-sample sizes like 2000 and 4000, with the underestimation bias being more visible for relatively smaller values of *h*^2^. The genetic relationship value between two individuals with British ancestry is usually very small (< 0.05). So, it is possible that for our small subset of individuals, the model became unidentifiable due to the GRM being close to the identity matrix. This would induce a large bias. However, the GREML (sub) estimates for large sub-sample size like 16,000 were nearly unbiased in all the cases, but had much larger variance compared to the corresponding PredLMM estimates.

#### 3.2.2 Simulation Study 2B: Simulation with a heterogeneous sub-population

Next, we studied the performance of the methods when the population is heterogeneous with multiple ethnicities. Every time we randomly selected 100,000 people from the mixed sub-population of size 157,000. Note that, unlike the last section, we did not choose people with only British ethnicity this time. Rest of the simulation procedure was the same as the earlier section. Box-plot comparison of the estimates are shown in Figure 4. GREML (sub) estimates this time did not show huge bias for small or moderate sub-sample sizes as they did for homogeneous population, but the estimates still showed very high variance. The issue of unidentifiability possibly did not arise here, because two individuals with different ethnicities often had high genetic relationship values (> 0.05). It kept the sub-sample GRM away from being close to the identity matrix. Similar to Simulation 2A, PredLMM showed slight downward bias when the true heritability was high (case (c)) and slight upward bias when the true heritability was low (case (a)). For moderate value of heritability (case (b)), the bias was negligible even for the smallest knot-size of 2000.

**Figure 4:**
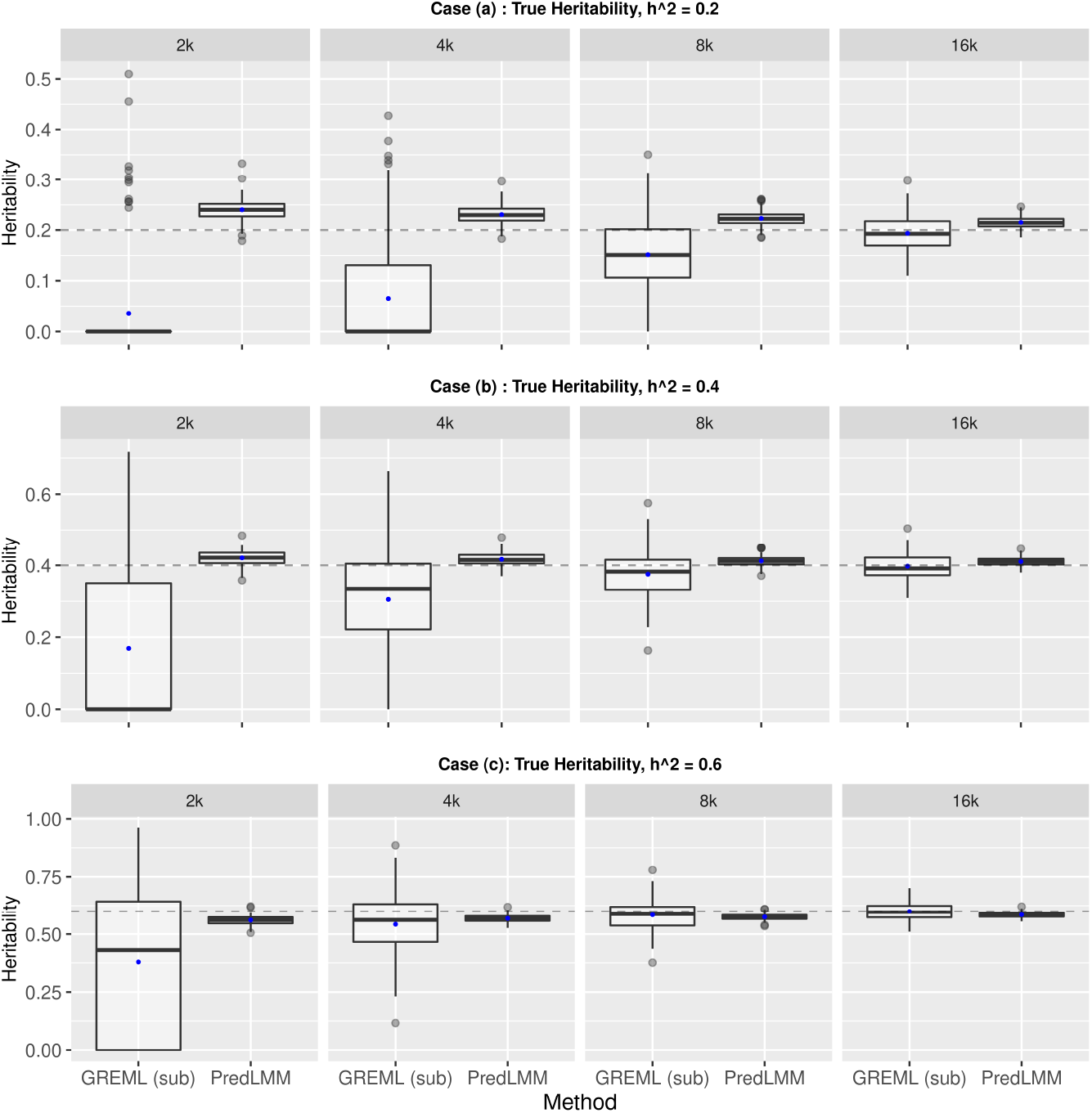
The figure shows boxplots of heritability estimates by GREML (sub) and PredLMM with four different sub-sample sizes: 2000, 4000, 8000, 16000 for cases (a), (b) and (c) from Simulation Study 2A (Section 3.2.1).

**Figure 5:**
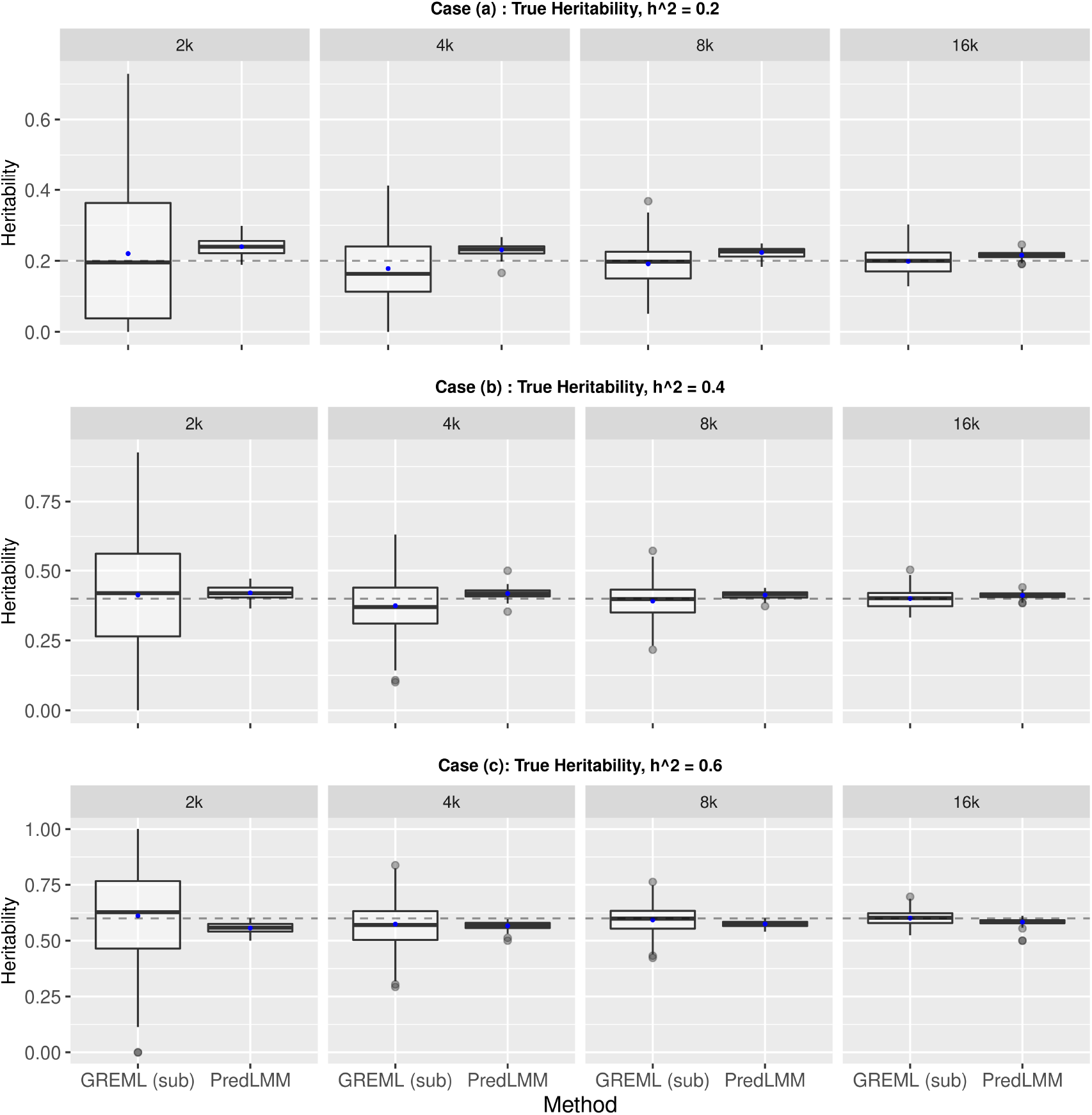
The figure shows boxplots of heritability estimates by GREML (sub) and PredLMM with four different sub-sample sizes: 2000, 4000, 8000, 160000 for case (a), (b) and (c) from Simulation Study 2B (Section 3.2.2).

### 3.3 Analysis of UK Biobank traits

We estimated heritability of a number of quantitative traits: *Standing Height, Weight, BMI, Diastolic and Systolic blood pressure, Hip and Waist circumference* using the British population of size 286,000 and 566,000 SNPs. We took into account the fixed effects of several covariates, such as sex, age, squared age, and the top 10 genetic principal components. We implemented the GREML (sub) and the PredLMM approaches with four different sub-sample (knot) sizes: 2,000, 10,000, 20,000 and 40,000. Since, running the full version of Bolt-REML would take an exorbitant amount of time, we calculated the approximate “pseudo-heritability” option in BOLT-REML [20, 22]. The results are displayed in Figure 6. For PredLMM and GREML (sub), we estimated heritability and its standard error for all these traits (*Standing Height, Weight, BMI, Diastolic and Systolic blood pressure, Hip and Waist circumference*), and plotted the point estimates along with 95% confidence interval (CI) in Figure 6. We truncated the CIs between 0 and 1. For Bolt-REML (Pseudo), however, we only plotted the point estimates since the package did not provide a standard error estimate.

**Figure 6:**
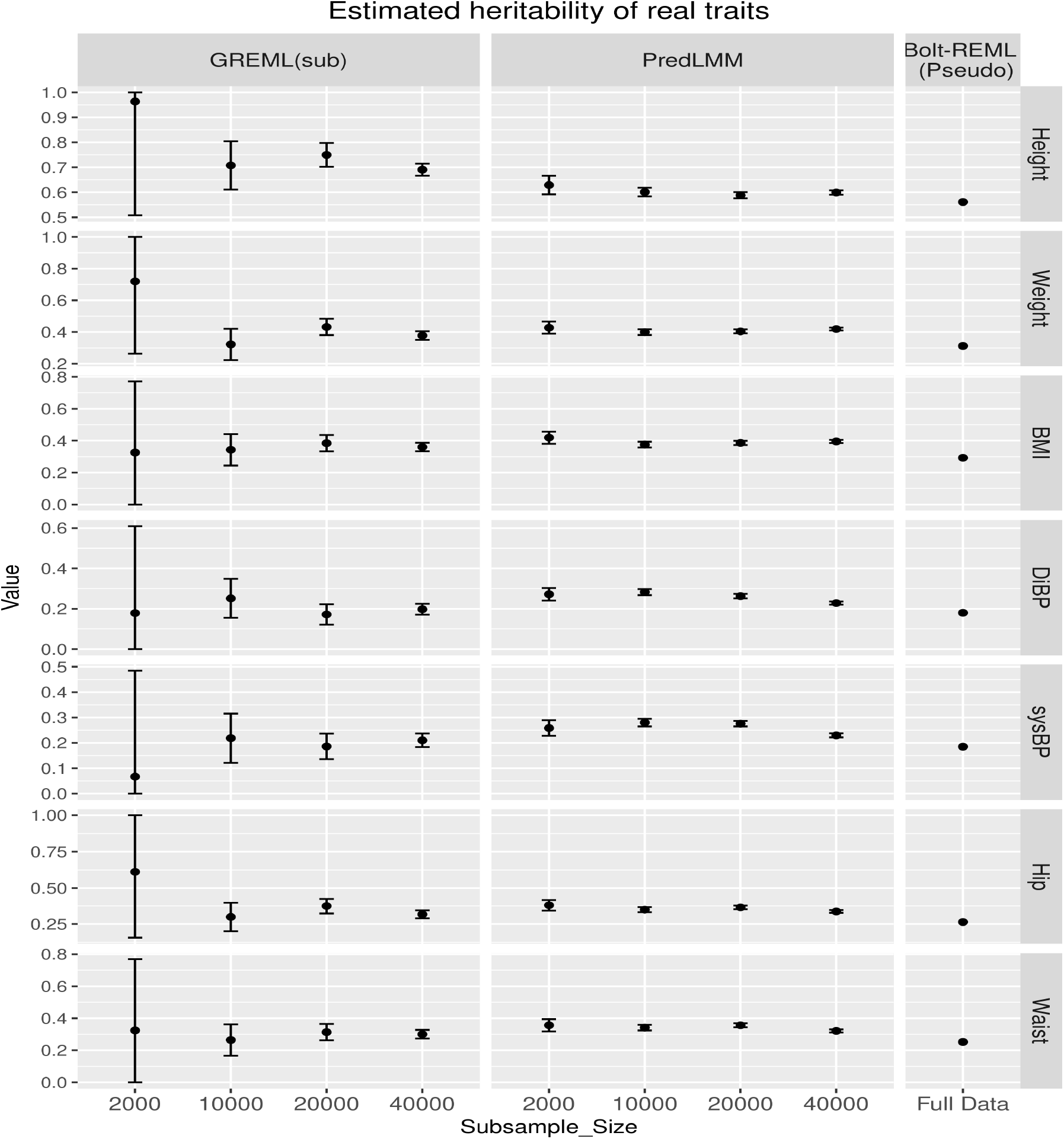
The figure shows the point estimates and 95% CI’s of the heritability estimates by PredLMM and GREML (sub) with varying sub-sample (knot) sizes and only point estimate by Bolt-REML (pseudo) for seven different real traits. The CIs’ have been truncated below at 0 and above at 1.

According to Figure 6, both PredLMM and GREML (sub) gave point estimates close to Bolt-REML (Pseudo) for higher sub-sample (knot) sizes. GREML (sub) estimates with the lowest sub-sample size 2000, were mostly unreliable. This was quite consistent with what we found in Simulation Study 1 in Section 3.2.1. It again validates our earlier conclusion that GREML (sub) can turn out to be highly unreliable if the sub-sample size is inadequate or the chosen sub-sample does not represent the population well. However, even with the smallest knot-size (2,000), PredLMM produced robust estimates of heritability with a narrow CI. For other knot-sizes, PredLMM estimates were very close to each other with the CIs becoming increasingly narrower. Overall, we found a slight decreasing trend in the PredLMM estimates as we increased the knot-size for majority of the traits. It may be similar to what we have seen in the simulation studies that PredLMM displays slight over-estimation with small knot-sizes if the true heritability is low. To conclude, just like the simulation setups, it is again observed that even with a very small number of knots relative to the total population size, PredLMM can produce highly precise and robust estimate of heritability.

## 4 Time Comparison

The huge time advantage of PredLMM has already been illustrated in Figure 2. Here, we present a few more tables in support of that and specify all the technical details. We ran all the methods on a HP Linux cluster with nodes that use 24 many Haswell E5-2680v3 processor cores. We have listed in Table 1 and Table 2 the time taken by different methods for Simulation 1 from Section 3.1 and for Simulation 2 from Section 3.2 respectively.

**Table 1:** Time comparison of different methods in seconds for Simulation Study 1 in Section 3.1 with 5k (8k SNPs) and 8k (13k SNPs) individuals.

|  | GCTA | GEMMA | Bolt-REML | GREML (500) | GREML (2000) | PredLMM (500) | PredLMM (2000) |
| --- | --- | --- | --- | --- | --- | --- | --- |
| 5k | 15.5 | 351.07 | 13.25 | 3.41 | 6.7 | 5.398 | 16.77 |
| 8k | 33.5 | 1293.44 | 27.87 | 5.7 | 3.3 | 13.67 | 28.46 |

**Table 2:** Time comparison (in minutes) of PredLMM for varying different knot (sub-sample) sizes with Bolt-REML for Simulation Study 2B in Section 3.2.1 with 100k individuals

|  | PredLMM |  |  |  | Bolt-REML |
| --- | --- | --- | --- | --- | --- |
| Knot size | 2000 | 4000 | 8000 | 16000 |  |
| Time | 28.33 | 31.41 | 44.17 | 80 | 986.4 |

From Table 1, we see that the methods like GCTA and Bolt-REML take similar amount of time, whereas PredLMM with 500 knots takes around 40% of that. PredLMM with 2000 knots takes time similar to Bolt-REML. The time advantage is more prominent in Table 2 which corresponds to the analysis from Simulation Study 2B (Section 3.2.2) with 100,000 individuals (this comparison is also shown in Figure 2).

According to Table 2, PredLMM takes just a fraction (around 8%) of time compared to Bolt-REML even if we choose a large knot size of 16,000. PredLMM takes very similar amount of time for knot sizes 2000 and 4000. We noticed a significant leap in the run-time from knot size of 8000 to knot size of 16,000. Recall that the per iteration computational complexity of PredLMM is *O*(*Nr*^2^ + *r*^3^) i.e the complexity is cubic with respect to the knot size *r* which justifies the leap. One may argue that it would be wise to use just 8000 knots since it can yield a reasonable estimate in a very reasonable time. We should also mention that we used a pre-computed GRM (using GCTA) in all our analyses (we computed the GRM for the entire population and used its sub-matrices as necessary in our Simulation Study 2). Computing the GRM is an arduous task that can take multiple hours depending upon the number of SNPs and the number of individuals. It has computational complexity of *O*(*MN* ^2^). But, it is usually of less concern since the computation is just a one time thing and the computed GRM then can be used in multiple analyses. Bolt-REML does not use a pre-computed GRM and uses the genetic data every time for each of the traits which makes it very time consuming to perform a heritability analysis with multiple traits.

## 5 Discussion

Genome-based restricted maximum likelihood (GREML) approaches for estimating heritability have become widely popular with the availability of large scale cohort studies. However, majority of the existing approaches such as GEMMA, GCTA, Bolt-REML implementing GREML, either become computationally very demanding or even intractable when the number of individuals (*N*) is too large. In this paper, we have developed a rapid algorithm for estimating heritability in large scale cohort studies. Our proposed approach PredLMM approximates the likelihood of a GREML approach. The approximation is achieved by unifying the concepts of genetic coalescence and Gaussian Predictive Process models. The algorithm reduces the usual per iteration computational complexity from *O*(*N* ^3^) to *O*(*Nr*^2^ + *r*^3^) where *r* (knot size) is much smaller than *N*.

From the simulation study of Section 3.1, we have seen that under the presence of genetic pattern (a tree like structure) among the individuals, PredLMM yields highly robust estimate of heritability even with a small knot size (*r*). To replicate more realistic scenarios, next we have performed simulation studies using the real genetic data from the UK Biobank cohort study. We have checked the performance of PredLMM in two cases, a highly homogeneous sub-population (see Section 3.2.1) and a heterogeneous sub-population (see Section 3.2.2) for a varied range of true heritability values. We have observed that even with a very small knot size (say 1% of the population size *N*), PredLMM can achieve an extremely reliable estimate of heritability at a fraction of time taken by existing softwares like Bolt-REML. PredLMM estimates also have much narrower CI’s compared to GREML (sub) which behave erratically if the sub-sample size is small. Finally, we have estimated the heritability of a number of quantitative traits like *Standing Height, Weight, BMI, Diastolic and Systolic blood pressure, Hip and Waist circumference* using the entirety of the British population from UK Biobank data. For all the traits, estimates by PredLMM for varying knot-sizes come out to be very close and also, very similar to other methods like GREML (sub) and Bolt-REML (Pseudo). Therefore, it further echoes our conclusion that with even a small set of knots PredLMM can yield a highly robust estimate of heritability in a fraction of time taken by other methods.

Our next goal would be to analyze behavioral traits like *Alcohol Consumption, CPD (cigarettes smoked per day)* etc. from the UK Biobank data. It would be slightly more challenging since those traits often tend to be skewed and deviate from the usual normality assumption. A very efficient module for implementing PredLMM in Python is available here, https://github.com/sealx017/PredLMM.. Also, so far we have used GCTA for computing the GRM (**A**) which takes up *O*(*N*^2^) storage and costs significant amount of time. PredLMM actually does not need computing of the full GRM but only a sub section of it requiring storage space of just *O*(*Nr* + *r*^2^). Therefore, in order to use less resources and further faster analysis of large scale data, we would like to incorporate the feature of computing just the required section of the GRM in our module as well.

## 6 Acknowledgement

This study is supported in part by the National Institutes of Health/National Institute on Drug Abuse grants 5R01DA033958-02 and 1R21DA046188-01A1.

## A Appendix

### A.1 Variance of PredLMM estimator

Here, we derive the variance of the PredLMM estimator. PredLMM considers the following multivariate normal distribution,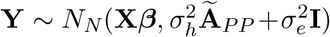. Lets denote the covariance matrix as, 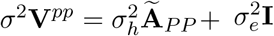 where 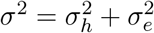 and **V**^*pp*^ is written as,

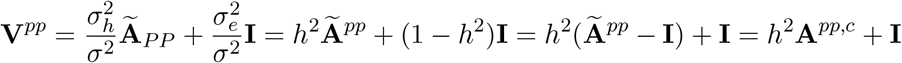

where **A**^*pp,c*^ = **Ã**^*pp*^ – **I** 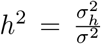 Lets denote the corresponding log likelihood by *l*^*pp*^ (**Y**). The first derivative *l*^*pp*^ (**Y**) w.r.t ***β*** *σ*^2^/*h*^2^ would be,

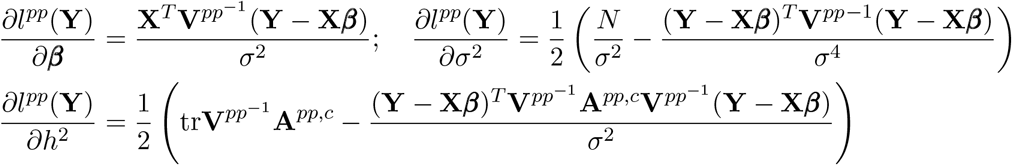

These three equations can be solved iteratively by Newtonm-Raphson method [33]. ***β*** and *σ*^2^ both will have closed form solutions at every iteration (*t*) obtained by equating the respective first derivatives to 0: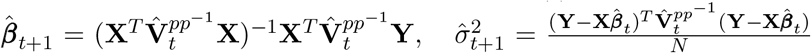 Where 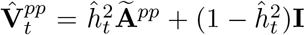 and 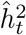 is the value of *h*^2^ at the *t*-th iteration.

The actual expression of the asymptotic variance of the estimator of *h*^2^ will involve the information matrices corresponding to parameters ***β***, *σ*^2^. We approximate the asymptotic variance by making a simplifying assumption. Upon estimating the values of ***β***, *σ*^2^, we treat them as known, thereby ignoring their distributional properties. The second derivative of *l*^*pp*^(**Y**) with respect to *h*^2^ would be,

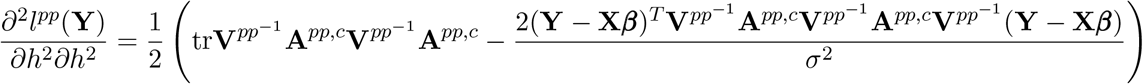

Next, we derive expectation of several important terms.

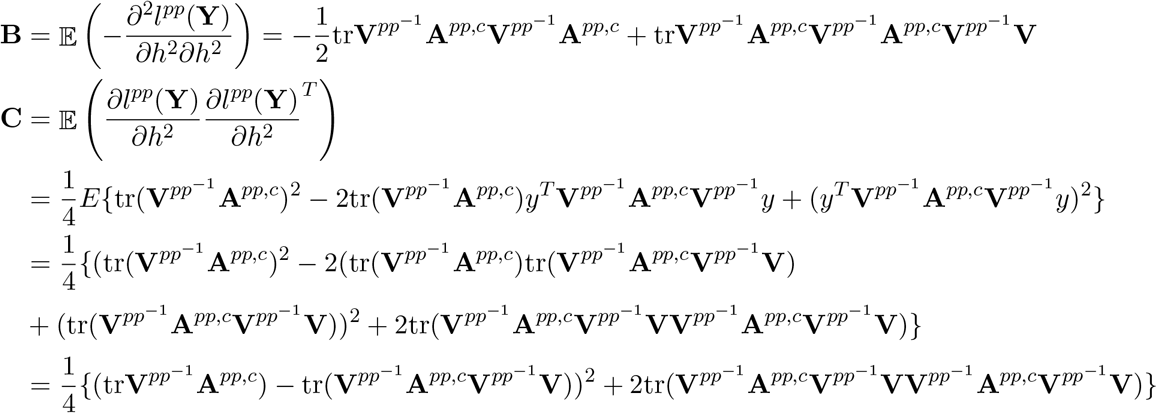

We have used the fact that the true distribution of **Y** is multivariate normal with mean **X*β*** and covariance matrix 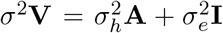 and hence, 𝔼 (**Y** − **X*β***)(**Y** − **X*β***)^*T*^ = *σ*^2^**V**. Then, the asymptotic variance of the PredLMM estimator would be **B**^−1^**CB**^−1^ [5]. Since, the expressions of **B** and **C** require dense matrix multiplications that can be computationally intractable, it is necessary to reasonably simplify them. We assume that **V**_*pp*_ ≈ **V** and it leads to, 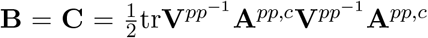. Finally, the variance formula reduces to,

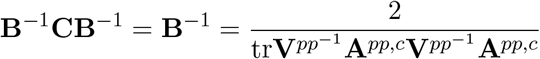

We have verified the validity of our variance estimate by looking at the coverage probabilities in the simulation studies.

